# Acceleration of vision changes object representations from infancy to adulthood

**DOI:** 10.64898/2026.10.02.756174

**Authors:** Céline Spriet, Emilie Serraille, Samar Naderi, Marwane Agrebi, Zied Bouyahia, Stéphane Derrode, Daniel Kaiser, Liuba Papeo, Jean-Rémy Hochmann

## Abstract

While much of the neural and cognitive machinery for visual perception is in place within weeks from birth, we identified and quantified a fundamental difference between infant and adult vision: the temporal rate at which meaningful visual information can be reliably extracted. Using frequency-tagging electroencephalography with rapid serial visual presentation, we found that object categorization at 4 and 9 months is respectively seven times and two to three times slower than in adults. Using quantitative information from infants’ experiments, we created and presented accelerated visual streams to adults, to match infants’ perceptual experience. We used deep convolutional neural networks to identify object representations that match those elicited in adults when accelerating the information flow. Results show that acceleration compromises finer-grained object distinction while sparing broad, general object representations. We propose that slow vision is a design feature of early development for preserving and highlighting the most stable and general features of the world.

## Introduction

Perception is the medium that connects the external world to our internal subjectivity, transforming sensory information into representations that can be used for reasoning, planning, acting, decision making and learning. The core neuronal and cognitive machinery for visual perception is largely in place early in life, possibly at birth (*1*–*3*). However, an immature brain and fledgling myelination (*4, 5*) impact the transmission and integration of information into complex representations (*6*–*11*), with consequences on the quality of young infants’ visual experience that remain unknown.

Contrary to an outdated belief (*12, 13*), infants are not “nearly blind”, nor are they unable to organize visual input into meaningful objects and scenes (*14*–*17*). Figure 1 illustrates what infants of different ages may experience when looking at a static colorful image. Visual acuity develops from about 0.5/10 at birth, to 1/10 at 4 months and 3/10 at 9 months (*18*). Neonates show spontaneous attraction towards faces and face-like patterns (*19*), segment objects from background on the basis of common motion (*20*) and recognize biological motion relying on patterns of semi-rigid motion (*21*). Neonates also discriminate fine orientations (*22*) and have size constancy (*23*). By 4 months, infants have developed sophisticated color vision (*24, 25*) and perceive depth through stereopsis (*26*).

**Figure 1.**
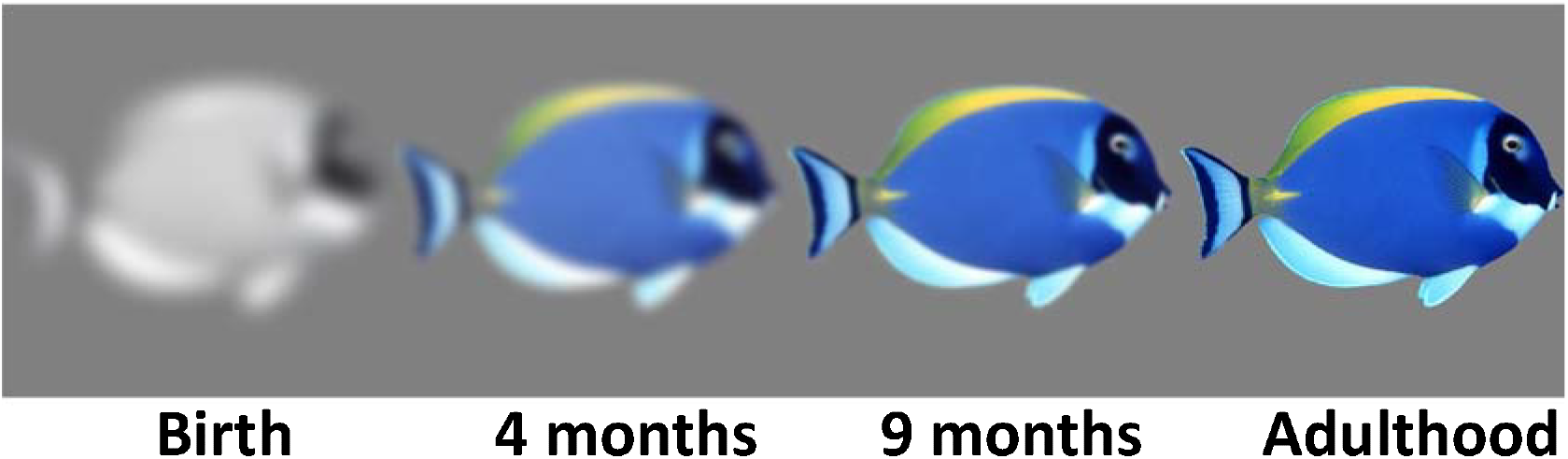
Illustration of infants’ vision. The four objects simulate the visual experience of static images at about 60 cm distance, as a function of visual acuity and color perception in newborns, 4- and 9-month-olds infants and adults, respectively. To simulate visual acuity, a Gaussian filter was applied to the original image (Adulthood). We used a 2-D Gaussian smoothing kernel with standard deviation of 20 for newborns, 10 for 4-month-olds and 3.33 for 9-month-olds, respectively corresponding to 0.5/10, 1/10 and 3/10 acuity (*18*) as compare to 10/10 acuity in adults. Color vision was simulated by setting a threshod on discriminable saturation. Threshold to simulate color vision at different ages were computed based on existing models (*24*). Original images were coded in the hsv format and the proper percentage of saturation was removed from the image: 88.9% for newborns, 24.4% for 4-month-olds and 11.1% for 9-month-olds.

But vision is more than the sensitivity to disconnected individual features like color, orientation, motion and texture, or to specific patterns that characterize biologically relevant stimuli such as faces. Mature vision binds features into object representations, e.g., roundness, yellowness and fuzzy texture, are integrated into a unified rounded yellow fuzzy object, that is, *a tennis ball* (*27*). The hierarchy of processing stages, from the analysis of basic visual features (roundedness, color, texture) to object representation (rounded yellow fuzzy object) lays at the bottom of object recognition and categorization (a tennis ball) (*14*). Broad categorical distinctions, such as those between animate and inanimate objects, can be extracted rapidly and are evident early in development, preceding the emergence of more fine-grained categorical representation (*28*). It has been suggested that young infants rely only on a subset of features available to adults (*29*) and prioritize salient information (*28*). As a result, their experience of the world may be quite different from that of older children and adults.

Prior work has largely focused on the quality of visual representations, with scant consideration of the *speed* of visual processing in infants. Yet, a common, everyday, observation is that young infants react slowly to sensory stimulations, as if they process information slowly. Relatively long delays, compared with adults, separate a loud noise from the infant’s headturn (*30*), an injection from their cry (*31, 32*), and their mom’s smile from their own smile. Recent studies have shown that neuronal intrinsic timescales are slower in neonates (*33*), and infants segment the visual world in fewer, longer events than adults (*34*–*36*). Previous work also suggests slower and more variable visually evoked responses during early life (*30, 37, 38*). Here, we provide the first empirical demonstration that infants are indeed slow, because of intrinsic limitations in their temporal processing capacity, i.e., the sampling rate at which meaningful visual information can be extracted from the continuous information input. In six experiments, we showed that, by 4 months, infants can categorize objects into animate *vs* inanimate, but this capacity is profoundly affected by the speed of visual stimulation: Infants encode incoming visual information at a pace that is significantly slower than adults. We will further argue that differences in sampling rate result in profound differences between infants and adults, in terms of informational content.

## Results

### Experiment 1-5 - Seeing animacy in infancy

#### Experiment 1

Visual objects can be categorized as *animate* or *inanimate*, leveraging visual differences that covary with categorical differences, including spectral properties, color, texture and shape (*39*–*43*). Here, to assess this process, we employed a large and diverse set of 640 animate and inanimate object images, most of which would be unfamiliar to young infants (see *Material and Methods* for a detailed description). Animacy categorization was tested with frequency-tagging electroencephalography (ftEEG; *44*–*47*). ftEEG measured the EEG responses to images presented in rapid sequence, at a regular *base* frequency, varying between 4 and 30 images per second (Figure 2A), with a regular category-change, such that every *n* examplars of a ‘standard’ category (e.g., inanimate) an image from another ‘oddball’ category (i.e., animate) was shown. In this paradigm, stimulus perception is signaled by a response in the EEG signal at the base frequency (i.e., 4 to 30 Hz; *Material and Methods*) and successful visual categorization is signaled by another distinctive response at the category-change frequency (Figure 2B). In adults, the amplitude of this category-change response correlates with explicit stimulus recognition (*43, 48, 49*).

**Figure 2.**
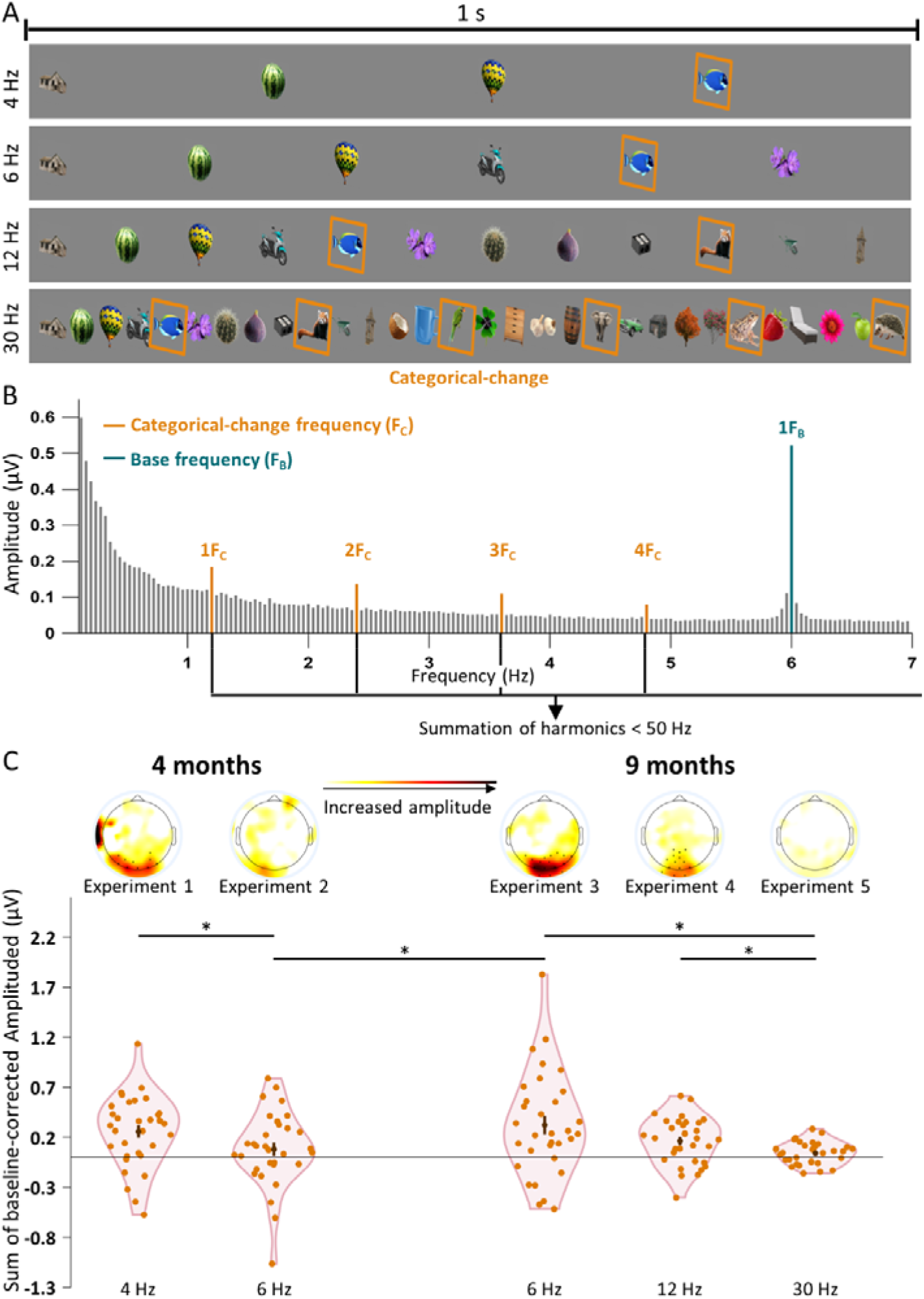
Experimental design and results of Experiments 1-5. **(A)** One-second extracts from a stream of visual stimuli testing the categorization of animates among inanimate objects at different base frequencies: 4, 6, 12 and 30 Hz. Images in orange squares are animate stimuli that appear rarely but regularly (category-change frequency). **(B) Example of the expected spectrum for a stimulation at F**_**B**_**=6 Hz with a category-change at F**_**c**_**=1.2 Hz**. Regular visual stimulation is expected to elicit visible peaks in the Fast Fourier Transform of the EEG signal. Response to the category-change is estimated by summing baseline-corrected amplitude at F_C_ and harmonics. Unspecific response to the visual stimulation is estimated by summing baseline-corrected amplitude at F_B_ and harmonics. **(C) Results of Experiments 1-5 with 4- and 9-month-olds**. Sum of baseline-corrected amplitudes for harmonics of F_C_ under 50 Hz, averaged over posterior electrodes showing a significant response for 4-month-olds with 4 Hz streams and for 9-month-olds with 6, 12 or 30 Hz streams. Dark dots on topoplots indicate the position of significant clusters of electrodes. Orange dots represent the response of each participant.

In Experiment 1, a group of 4-month-old infants (N=32) was tested with streams of images presented at a frequency of 4 Hz, with one category-change every 4 images (category-change frequency: 4/4 = 1 Hz; video S1). A one-sided one-sample *t*-test over posterior electrodes revealed a significant category-change response (*M*_*Amplitude*_±*sd* = 0.257±0.356; 95% CI = 0.151 – Inf; *t*(31) = 4.085; *P* = .0001; *d* = 0.722; Figures 2C, 3A). A cluster mass permutation test in these posterior electrodes further circumscribed the effect to a cluster of ten posterior electrodes (Figures 2C, 3A) (*M*_*Amplitude*_±*sd* = 0.363±0.427; *P* = .001). In a control group (Experiment S1, *Supplementary material;* video S2), we showed that this response was absent for phase-scrambled images (over posterior electrodes: *M*_*Amplitude*_±*sd* = 0.075±0.435; 95% CI = −0.056 – Inf; *t*(31) = 0.969; *P* = .170; *d* = 0.171; no significant cluster of electrodes), and significantly higher for original images compare to phase-scrambled images (*M*_*Difference*_±*sd* = 0.183±0.397; 95% CI = 0.017 – Inf; *t*(62) = 1.841; *P* = .035; *d* = 0.460; Figure 3A).

**Figure 3.**
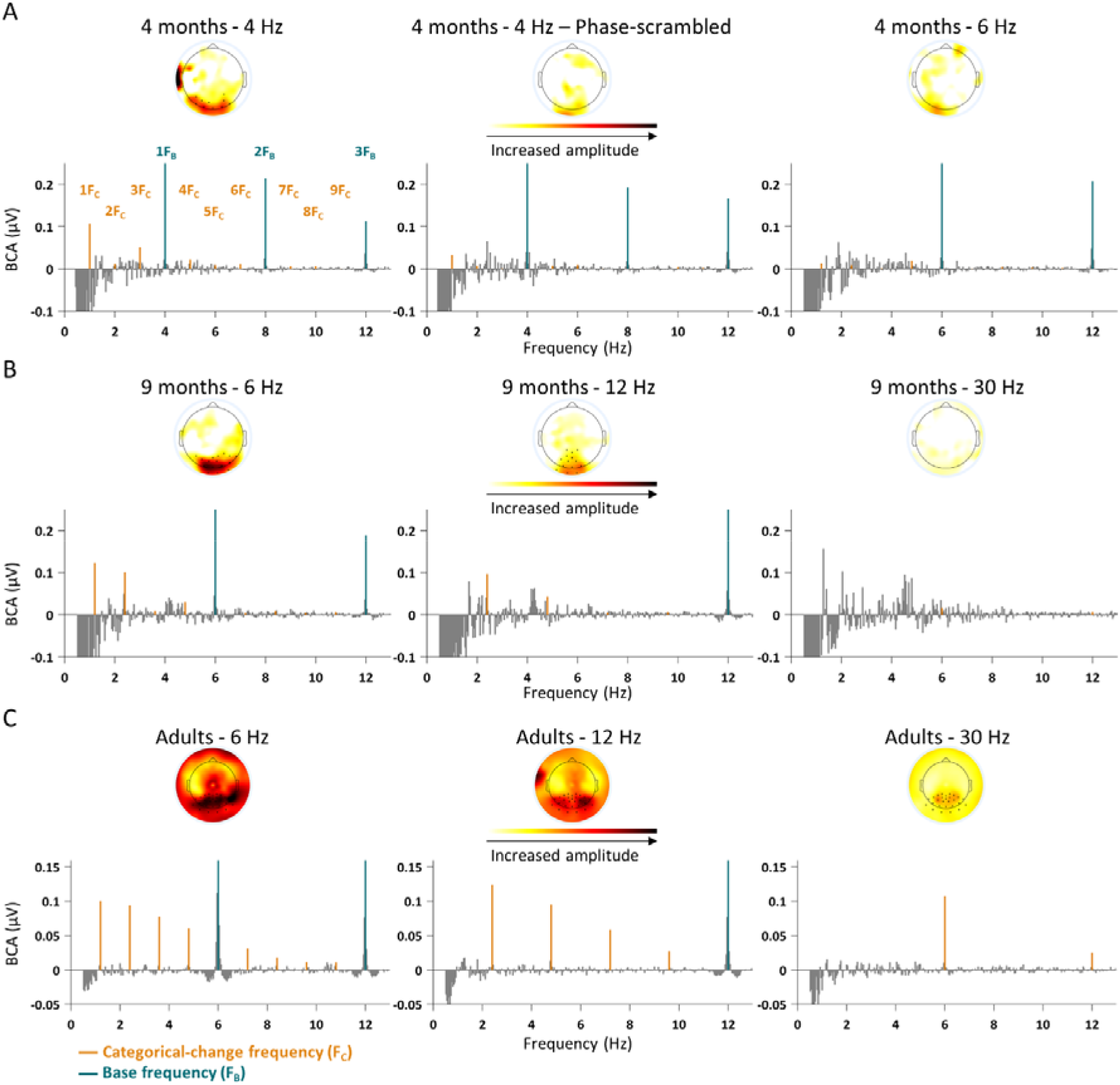
Baseline-subtracted amplitudes for all experiments. Results of Experiments 1, S1 and 2 (A), Experiments 3-5(B) and Experiment 6 (C). Baseline-corrected amplitudes (BCA), averaged over posterior electrodes and all participants of each group. Baseline-corrected amplitude responses were computed by subtracting the average amplitude of surrounding frequencies from the amplitude of the frequency. Dark dots in the topoplots represent the electrodes showing a significant response for the group. Orange lines represent the harmonics of the categorical-change frequency; blue lines represent the harmonics of the base frequency.

Because phase-scrambled images match the original images for low-level features such as surface area, spatial frequency distribution, and luminance, color and contrast variations, Experiment S1 rules-out that the effect observed in our ftEEG paradigm could be fully explained by systematic differences in low-level features between animate and inanimate objects. Rather, the effect signals the extraction of complex information diagnostic of the animate/inanimate categories.

#### Experiment 2

In Experiment 2, we tested a new group of 4-month-olds (N=32) with accelerated streams of stimuli presented at 6 Hz and involving a category-change every 5 stimuli (category-change frequency: 6/5 = 1.2 Hz; video S3). A one-sample *t*-test one-sided over posterior electrodes revealed no significant category-change response (*M*_*Amplitude*_±*sd* = 0.081±0.373; 95% CI = −0.031 – Inf; *t*(31) = 1.226; *P* = .115; *d* = 0.217). Moreover, no significant cluster was found among these electrodes (Figures 2C, 3A). The category-change response in posterior electrodes was significantly higher in the group tested with 4 Hz-streams (Experiment 1) than in the group tested with 6-Hz streams (Experiment 2) (*M*_*Difference*_±*sd* = 0.177±0.365; 95% CI = 0.024 – Inf; *t*(62) = 1.936; *P* = .029; *d* = 0.484; Figure 2C). In sum, 4-month-olds can discriminate between animate and inanimate objects. However, increasing the rate of stimulus presentation from 4 to 6 Hz dramatically reduces, indeed abolishes, the neural signature of object categorization.

#### Experiment 3

In Experiments 3-5, we asked whether older infants could succeed where the younger failed, and in even more extreme conditions. Experiment 3 tested a group of 9-month-olds (N=32) with streams of images presented at a base frequency of 6 Hz with a category-change every 5 stimuli (category-change frequency: 6/5 = 1.2 Hz; video S3). A one-sided one-sample *t*-test over posterior electrodes revealed a significant category-change response (*M*_*Amplitude*_±*sd* = 0.322±0.517; 95% CI = 0.167 – Inf; *t*(31) = 3.522; *P* = .0007; *d* = 0.623; Figures 2C, 3B). This response was significantly larger than the response of 4-month-olds (Experiment 2) to the same 6-Hz streams (*M*_*Difference*_±*sd* = 0.241±0.451; 95% CI = 0.053 – Inf; *t*(62) = 2.140; *P* = .018; *d* = 0.535). A cluster-mass permutation test further circumscribed the effect to a cluster of sixteen posterior electrodes (*M*_*Amplitude*_±*sd* = 0.449±0.557; *P* = .0006; Figures 2C, 3B).

#### Experiments 4-5

In Experiments 4-5, we tested two new groups of 9-month-olds (N=32 each). In Experiment 4, streams of images were presented twice as fast as in Experiment 3 (base frequency: 12 Hz) with a categorical change every 5 stimuli (category-change frequency: 12/5 = 2.4 Hz; video S4). In Experiment 5, streams were shown at the extremely high frequency of 30 Hz with a categorical change every 5 images (category-change frequency: 30/5 = 6 Hz; video S5). Results showed that, in this age group, a category-change response persisted in posterior electrodes for both groups, but it was weaker with faster presentation rates (see below) (Experiment 4: *M*_*Amplitude*_±*sd* = 0.162±0.235; 95% CI = 0.091 – Inf; *t*(31) = 3.898; *P* = .0002; *d* = 0.689; Experiment 5: *M*_*Amplitude*_±*sd* = 0.041±0.112; 95% CI = 0.008 – Inf; *t*(31) = 2.091; *P* = .022; *d* = 0.370). Cluster-mass permutation tests identified a posterior cluster of 14 electrodes with 12 Hz streams (*M*_*Amplitude*_±*sd* = 0.257±0.241; *P* = .0002) but no significant cluster with 30-Hz streams (*P* > .10) (Figures 2C, 3B).

Statistical comparisons between the three groups of 9-month-olds tested with 6-, 12- and 30-Hz streams respectively (Experiment 3-5) showed significant differences (one-way ANOVA; *F*(2,95) = 5.683; *P* = .005; η^2^ = 0.109; Figure 2C). The category-change responses for 6-Hz streams and 12-Hz streams were comparable (*M*_*Difference*_±*sd* = 0.160±0.402; 95% CI = −0.008 – Inf; *t*(62) = 1.595; *P* = .058; *d* = 0.399), and significantly higher than the response for 30-Hz streams (6-Hz *vs*. 30-Hz: *M*_*Difference*_±*sd* = 0.281±0.374; 95% CI = 0.125 – Inf; *t*(62) = 3.002; *P* = .0021; *d* = 0.751; 12-Hz *vs*. 30-Hz: *M*_*Difference*_±*sd* = 0.121±0.184; 95% CI = 0.044 – Inf; *t*(62) = 2.625; *P* = .006; *d* = 0.656). Thus, acceleration from 6 to 12 Hz had no significant impact on object categorization in 9-month-olds, whereas acceleration to 30 Hz disrupts it. Furthermore, we observed that acceleration of image presentation also affected the response at the base frequency. However, this effect showed a different pattern, with a large decrease from 6 to 12 Hz (*Supplementary Material*). This dissociation indicates that the modulation of the category-change response is not simply a spillover of sensory modulation, but reflects the contribution of higher-level processes.

In summary, 4-month-olds only showed a category-change response when images were presented at the relatively slow rate of 4 Hz (Experiment 1), but not when the rate accelerated to 6 Hz (Experiment 2). Nine-month-olds showed a category-change response with images presented at 6 Hz and up to 12 Hz (Experiments 3-4); the response was significanlty recuced, almost abolished, at 30 Hz (Experiment 5). These results demonstrate that the infant visual perception represents the animate and inanimate distinction already at the very young age of 4 months. The limits of this capacity depend on the stimulus presentation rate. Since successful categorization with faster stimulus presentation implies faster extraction of relevant visual information, these results demonstrate an acceleration of the information processing speed by a factor of about 3 between 4 and 9 months.

### Experiment 6 - Seeing like infants: The impact of processing speed on object perception

In Experiment 6, we asked how the information processing speed affects object representation, specifically, which information is preserved and which is lost when the rate of visual stimulation exceeds information processing limits. To test this, three groups of adults (N=12 per group) were tested with streams of objects presented at 6, 12 or 30 Hz, respectively, with a category-change every five images (category-change frequencies: 6/5 = 1.2 Hz; 12/5 = 2.4 Hz; 30/5 = 6 Hz). All groups exhibited a significant category-change response over posterior electrodes (one-sample *t*-test one-sided; 6 Hz group: *M*_*Amplitude*_±*sd* = 0.420±0.227; 95% CI = 0.302 – Inf; *t*(11) = 6.397; *P* < .0001; *d* = 1.847; 12 Hz group: *M*_*Amplitude*_±*sd* = 0.321±0.147; 95% CI = 0.245 – Inf; *t*(11) = 7.573; *P* < .0001; *d* = 2.186; 30 Hz group: *M*_*Amplitude*_±*sd* = 0.154±0.113; 95% CI = 0.095 – Inf; *t*(11) = 4.724; *P* = .0003; *d* = 1.364). Cluster-mass permutation analyses confirmed the effect in all posterior clusters (6 Hz group: *P* = .0004; 12 Hz group: *P* = .0004; 30 Hz group: *P* = .0006) indicating that object categorization was preserved even with 30-Hz streams. The response nevertheless varied with stimulus presentation rate (one-way ANOVA on posterior electrodes: *F*(2,35) = 7.599; *P* = .002; η^2^ = 0.315): it was comparable for 6- and 12-Hz streams (*M*_*Difference*_±*sd* = 0.099±0.191; 95% CI = −0.035 – Inf; *t*(22) = 1.269; *P* = .109; *d* = 0.518), but significantly reduced for 30-Hz streams (6 Hz vs 30 Hz streams: *M*_*Difference*_±*sd* = 0.267±0.179; 95% CI = 0.141 – Inf; *t*(22) = 3.638; *P* = .0007; *d* = 1.485; 12 Hz vs 30 Hz streams: *M*_*Difference*_±*sd* = 0.167±0.131; 95% CI = 0.076 – Inf; *t*(22) = 3.134; *P* = .002; *d* = 1.280) (Figures 4A, 3C). In sum, as the stimulus presentation rate increased from 12 to 30 Hz, the category-change response indexing visual object categorization diminished, although it was not abolished. Importantly, the modulation of the category-change response followed a different pattern than the response at the base frequency (*Supplementary Material*), suggesting the category-change response reflects the contribution of high-level processes.

**Figure 4.**
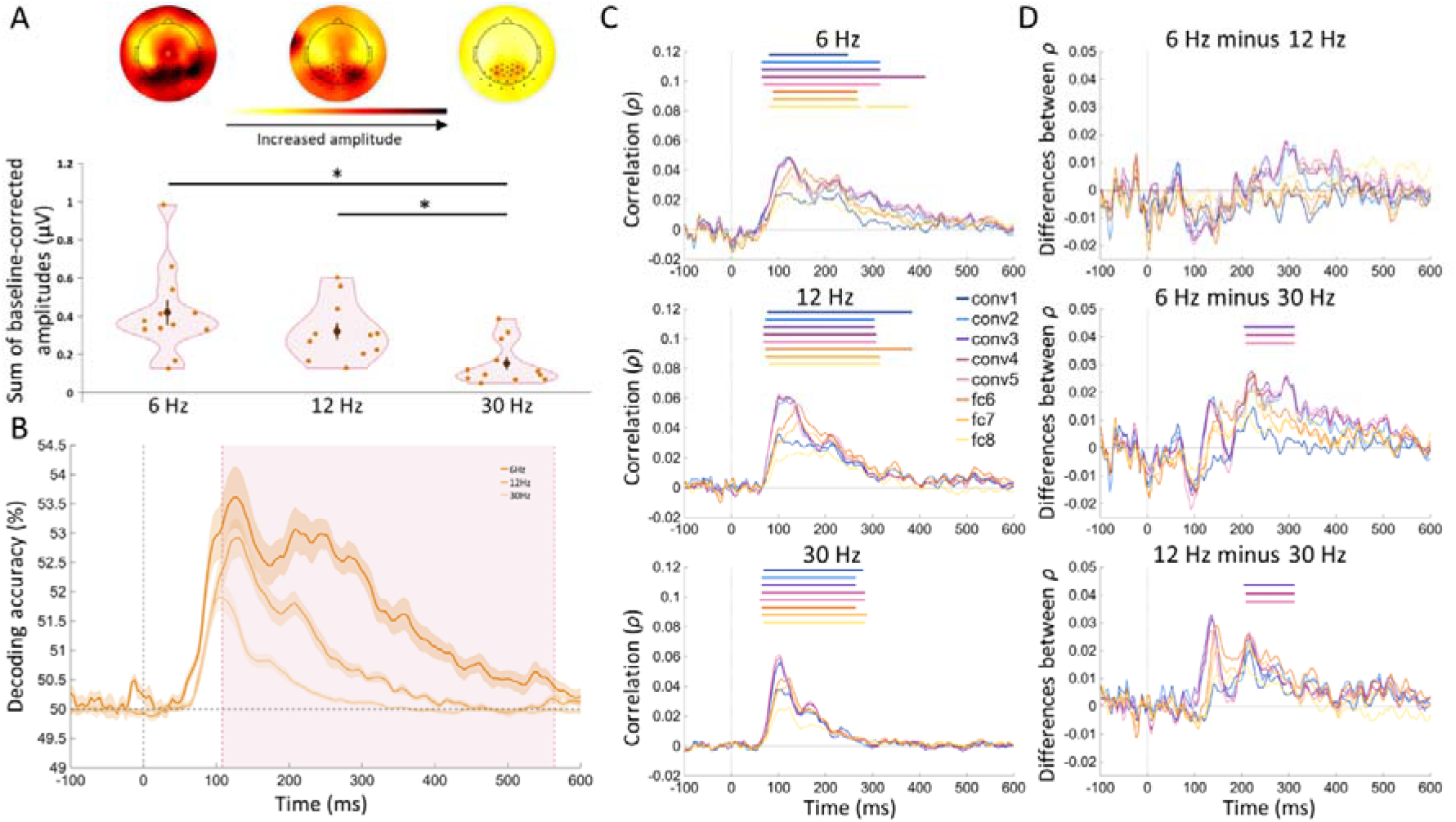
Results of Experiments 6 with adults. **(A)** Sum of baseline-corrected amplitudes for harmonics of F_C_ under 50 Hz, averaged over posterior electrodes showing a difference between groups for each age. Dark dots in the topoplot represent the electrodes showing a significant effect at the group-level. Orange dots represent the response of each participant. **(B-D) Decoding accuracy and correlation with CNNs. (B)** Grand average decoding accuracy between all pairs of objects, for each group in Experiment 6. The orange area indicates the time window revealing a significant difference between groups (cluster-mass permutation ANOVA analysis). **(C)** Results of the RSA between the EEG decoding and the 8 layers of AlexNet at each time point of the EEG signal. Significant correlations are denoted by colored bars for each layer. **(D)** Differences between the correlation of each group and layer of AlexNet. Significant differences between groups are denoted by colored bar for each layer.

### Characterizing the impact of the stimulus presentation rate on object representations

The frequency tagging design focused on the broad animate-inanimate categorization. However, adults can also represent objects at basic and subordinate category levels (e.g., an animal as a dog or a Dalmatian). Narrower categories (dog or Dalmatian) rely on the availability and integration of more features than the animate/inanimate distinction. We reasoned that, when information influx exceeds processing capacity, some features may go unattended, with an impact on the ability to represent narrower/finer categorical distinctions. On this reasoning, increasing the stimulus presentation rate should reduce the breadth of features encoded for an object, thereby limiting the distinctions that can be made between objects. The following analysis confirmed this prediction, providing a model of what (which features) may be lost vs. spared when infants are exposed to stimuli that change faster than their sampling rate capacity.

First, for each object-image and time point, we extracted a multivariate EEG pattern. Stimulus identity discriminability was tested by measuring the classification accuracy for each of 102080 pairs of stimuli with multivariate pattern analysis (MVPA). Reliable object discrimination (i.e., above-chance decoding accuracy) was found at all three stimulus presentation rates (6, 12 and 30 Hz) with two early peaks (around 120 ms and 208 ms), compatible with the latency of object processing in visual cortex (*50, 51*). However, a cluster-mass permutation test showed that decoding accuracy steadily decreased as stimulus presentation rate increased, between 108 ms and 564 ms (*P* = .0002; Table S1; Figure 4B)^1^.

Second, pairwise classification accuracies were entered into a dissimilarity matrix modeling object representations in our adult participants, separately for each stimulation rate. To approximate hierarchical feature representations across time, we used AlexNet, an eight-layer artificial deep convolutional neural network (CNN) trained on image classification (*52*). From layer-wise CNN activations, we created eight dissimilarity matrices by computing the similarity (1-ρ) of activation vectors for each stimulus pair. Finally, using representational similarity analysis (RSA, see *Material and Methods*), we measured to what extent object representation in the adults’ brain (Experiment 6) aligned with object representations across the eight layers of AlexNet, as visual stimulation rate increased.

Human object representation at all three visual stimulation rates correlated with object representation across all layers of AlexNet, with two peaks aligning with the decoding accuracy peaks (Table S2; Figure 4C). The acceleration of visual stimulation, especially from 12 to 30 Hz, reduced correlations of neural representation with mid-level layers *conv3, conv4* and *conv5* of AlexNet in time windows corresponding to the second peak or later, 204-312 ms (cluster-mass permutation tests; Table S3; Figure 4D). The same analyses with other CNNs (VGG19, Cornet-RT, Cornet-S, Cornet-R) yielded similar results (*Supplementary material*).

These results suggest that very rapid (30 Hz) visual stimulation preserves low-complexity features, accessed early (<204 ms) and encoded in early visual areas, modelled by early layers of the CNN (*conv1* and *conv2*). However, it compromises the encoding of higher-complexity visual features computed later and in more anterior visual areas, modelled by mid-level layers (*conv3, conv4* and *conv5*). These results show that when the temporal processing capacity of the visual system is exceeded, mid-level feature representations deteriorate. This pattern offers novel insight into infants’ visual perception. Particularly, fast changes in the visual environment, including those that leave adults’ obejct categorization unhindered, may limit the infants’ ability to access (or encode) mid-level features, biasing infants’ object categorization towards broader distinctions.

## Discussion

Here, we assessed the speed of visual object processing in infancy and how an information influx that exceeds temporal processing capacities affects object representation. Our study yielded four key results.

First, infants as young as 4 months can represent the broad animate-inanimate categorical object distinction in a wide, heterogeneous set of largely unfamiliar objects. In fact, our stimuli included hundreds of images, depicting mainly animals and objects that western city infants normally do not encounter (e.g., we can confidently assume none of the 4-month-olds tested here has ever encountered elephants, bears, cheetahs, volcanoes or hot-air balloons). This suggests that young infants can assign category membership on the first encounter with a stimulus, on the sole basis of brief static visual information.

Second, in the first months of life, infants’ perception is slow: 4-month-olds require visual presentation rates at least seven times slower than adults to extract information relevant for categorical (animate-inanimate) distinctions (4 vs. 30 Hz). This suggests that 4Hz streams may appear to 4-month-olds much like 30 Hz streams appear to adults (Videos S6-S8 vary playback speed to simulate vision at different ages). Taken together with the results of Experiment 6, the overall pattern suggests that, when presented with the same visual stream, infants extract substantially less information than adults. Some features may therefore go unattended and remain unavailable to infants, limiting their ability to represent fine-grained distinctions between objects. At first sight, slow perception, with its consequent reduction in the amount of information extracted and more limited categorization, might appear as a disadvantage. However, slow perception may constitute a design feature of early development, serving as an adaptive filter that prioritizes more stable and enduring properties of the environment over rapidly changing aspects, and fewer, but broader categories over fine-grained ones (*53*). This may facilitate generalization of knowledge, such that whatever is learned about, say, a cat can be generalized to virtually all animals or objects with featural similarities (to the superordinate level *animal* or *mammal* rather than at the basic level *cat*). In this vein, slow perception would be a case for the “less is more” hypothesis that highlights the potential benefits of limited cognitive functions and immature perception in early development (*54*–*56*): In sum, by starting slow, perception starts broad, highlighting the most stable and general features of the world.

Third, we simulated the effects of a visual stimulation that exceeded temporal processing capacity in adults. Using CNNs, we observed a deterioration of mid-level feature representations that typically emerge during sustained perceptual processing around 200-300 ms post stimulus-onset. This effect is analogous to the effects of visual masking, where, by interrupting recurrent processing, backward masking reduced the representation of mid-to high-level features in object-selective cortex, in a similar time window (*57*). This parallel suggests that, analogously to backward masking, the acceleration of visual stimulation disrupts recurrent processing.

Fourth, vision accelerates quickly in the first year of life: at 12 Hz, 9-month-olds see what 4-month-olds only see at 4 Hz. As infants grow, acceleration of perception would allow infants to detect and represent more features of the objects, and therefore to represent finer-grained distinctions between them, giving rise to new, more refined, categories (*28*). Dramatic changes between 4 and 9 months occur for many other functions. Four-month-olds lay prone or supine and depend on adults to be moved around, while 9-month-olds can sit and begin crawling (*58*–*60*). Four-month-olds produce cooing vocalizations, while 9-month-olds have already begun to produce language-like articulated speech (*61*). Nine-month-olds also reach a major milestone in social cognition with the emergence of joint attention (*62*). All these changes suggest that, between 4 and 9 months of age, infants become increasingly active and interactive, requiring finer tuning and faster adaptation to the environment. Cotemporality invites speculations about a relationship between these aspects of development. Building on the present work, future research may ask whether self-locomotion triggers the acceleration of perception or whether both result from a third factor, and the degree to which acceleration of information processing contributes to the development of language and social cognition.

In conclusion, the present work opens a breach in the mystery of how young infants experience the world. They may experience a fast-moving impoverished visual environment. We propose that infants’ perception acts as a filter that supports the formation of broader category representations, facilitating generalization, and therefore learning at the youngest age (*63*–*65*). This type of data selection mechanism may inspire the development of artificial intelligence that is less data-hungry and more parsimonious (*54, 56, 66*)–just like a learning human infant. Finally, our work reveals the limits of infants’ vision in a fast-paced world that is designed by and for adults. The quantitative characterization of infants’ visual perception capacities, relative to those of adults may represent a first step toward developing evidence-based reccomendations for designing more infant-friendly environments.

### Material and methods

#### Participants

Experiment 1 involved 32 4-month-old infants tested with 4 Hz streams (19 males; mean age 139.2±10.1 days). Two additional infants were tested in that Experiment but excluded from the analysis due to an insufficient number of trials included (see preprocessing), or crying and moving too much during the experiment.

Experiment 2 involved 32 4-month-old infants tested with 6 Hz streams (17 males; mean age 137.1±8.5 days). Five additional infants were tested in that Experiment but excluded from the analysis due to an insufficient number of trials included (see preprocessing), or crying and moving too much during the experiment.

Experiment 3 involved 32 9-month-old infants tested with 6 Hz streams (12 males; mean age 287.9±9.3 days). Five additional infants were tested in that Experiment but excluded from the analysis due to an insufficient number of trials included (see preprocessing), or crying and moving too much during the experiment.

Experiment 4 involved 32 9-month-old infants tested with 12 Hz streams (21 males; mean age 287.7±8.3 days). One additional infant was tested in that Experiment but excluded from the analysis due to an insufficient number of trials included (see preprocessing), or crying and moving too much during the experiment.

Experiment 5 involved 32 9-month-old infants tested with 30 Hz streams (12 males; mean age 292.3±19.3 days). Three additional infants were tested but excluded from the analysis due to an insufficient number of trials included (see preprocessing), or crying and moving too much during the experiment.

The sample size for infants’ experiments (N=32) was chosen based on a statistical power analysis performed on prior work using frequency-tagging to study the categorization of human faces (*44*). In that study, 15 4-month-old infants showed evidence of categorizing face stimuli. The minimal sample size for the observed effect was estimated to be 9 (Cohen’s *d* = 1.298, power = 0.95, α = 0.05; GPower 3.1). In the present study, we expected a smaller effect size as we were studying categories with higher within-category variability than the human face category. Thus, we decided to test 32 infants per group.

After having collected the full sample for Experiment 1, we verified that this sample size was larger than the minimal sample size (N=23) required to obtain the effect (amplitude higher than noise-level averaged over posterior electrodes: *M*_*Amplitude*_±*sd* = 0.257±0.356; 95% CI = 0.151 – Inf; *t*(31) = 4.085; *P* = .0001; *d* = 0.722; power = 0.95, α = 0.05; GPower 3.1).

Experiment 6 involved 36 adults divided into 3 groups of 12 adults each. One group was tested with 6 Hz streams (8 males; mean age 26.2±5.0 years). The second group was tested with 12 Hz streams (3 males; mean age 28.3±6.9 years). The third group was tested with 30 Hz streams (4 males; mean age 28.5±7.5 years).

The sample size for the adults’ experiment (N=12) was chosen following a power analysis of the results of an experiment with a similar design (*43*).

#### Stimuli

Stimuli were 640 colored photographs of isolated real-world objects picked from the internet, with the intention to reflect as much as possible the actual variability of each category in the real world. The animate set included 51 fish (from goldfish to sharks, including flying fish), 70 birds (from hummingbirds to emus, including various birds of prey), 179 nonhuman mammals (from mice to elephants including 10 primates, 5 cats and 8 dogs), 14 amphibians and 6 turtles, all different from one another. Humans were excluded to prevent a bias in the animate-inanimate categorization. Insects, spiders and reptiles (except for turtles) were excluded to prevent emotional reactions (e.g., disgust or fear). The inanimate set included 223 artificial objects (16 exemplars of buildings and constructions, 108 exemplars of clothes, pieces of jewelry, buttons, coins, and tools, 66 pieces of furniture and 33 vehicles) and 97 natural objects (54 different fruits and vegetables, and 43 different flowers, bushes and trees) of large (41%; e.g., planes, trees) or small (59%; e.g., cups, fruits) real-world size. All stimuli were superimposed on a gray background, and each image was resized to 629×629 pixels.

All the stimuli are available https://osf.io/jsvh4.

#### Experimental procedure

Participants sat on a chair (infant participants sat on their parent’s lap) ~60 cm away from a 60 Hz computer screen (resolution 1920×1200 pixels, size 51.5×32.2cm), where stimuli were presented centrally (16° of visual angle). Stimuli were presented via fast periodic visual stimulation, at the frequency of 4, 6, 12 or 30 Hz (250, 166.67, 83.33 or 33.33 ms per image), using Psychtoolbox (*67*) and MATLAB (The MathWorks, Nantucket, MA) in a squarewave design.

#### Experiments 1-5

Each infant was tested in one condition only, defined by the base frequancy of the sequence of images (4, 6, 12 or 30 Hz). Each infant took a maximum of 16 trials. Each trial lasted 32s and consisted of one stream of images, pseudo-randomly selected from a standard category (animate or inanimate) with the regular insertion, every four (4 Hz streams) or five (6, 12 and 30 Hz streams) images, of one image of the other category, leading a category-change stimulation of 4/4 = 1 Hz; 6/5 = 1.2 Hz; 12/5 = 2.4 Hz or 30/5 = 6 Hz (Figure 2 A). Four-month-olds were tested with 4 Hz or 6 Hz streams (Experiments 1-2), and 9-month-olds with 6, 12 or 30 Hz streams (Experiments 3-5). An infant only saw trials that all shared the same category-change (either animate or inanimate). Each trial started with a fade-in (increase in contrast) of 2 s and ended with a fade-out (decrease in contrast) of 2 s, to ease the presentation for the participant’s eyes and avoid eye movement caused by an abrupt appearance or disappearance (*44*). The experiment ended when all the 16 trials were shown, or when the infant started to cry, fell asleep, moved a lot or stopped looking at the screen anymore.

#### Experiment 6

The procedure was identical to that of Experiments 1-5, except for the following aspects. Participants were adults, and each was tested with 32 trials. Half of the trials used animate as the category-change and inanimate as standard, and half used inanimate as category-change and animate as standard, so that one adult participant saw both types of trials. Second, at the end of each trial, participants were presented with one image from the standard category (*e*.*g*., animate) to avoid creating an attentional bias toward the category-change (*e*.*g*., inanimate), as attention could enhance categorization (*68*). Participants were asked to say if they saw the image in the previous stream, by pressing one of two keys. This task was only designed to help participants focus on the screen and the experiment; these data were not analyzed.

#### EEG recording

EEG data were acquired using EGI amplifier GES 400 and 128-channel EGI geodesic nets. Data were acquired with vertex reference, using the EGI Net Station acquisition software, sampled at 1kHz. Impedance was lowered for each participant below 40kΩ. During the experiment, triggers were sent from the experimental computer to the acquisition one via a light sensor, allowing high timing precision.

### FT-EEG

#### Preprocessing

All preprocessing was performed using EEGLAB toolbox (*69*) and MATLAB R2023b (MathWorks). Raw data for each participant were first filtered using a 4th-order high-pass butterworth filter at 0.1 Hz and a 4th-order low-pass butterworth filter at 100 Hz. When needed, electrodes were interpolated (no more than 3 electrodes per participant). For adult participants, all electrodes were re-referenced using the average of all electrodes as a reference. For infants, we removed outer band electrodes due to the presence of increased noise (i.e., electrodes 1, 8, 14, 17, 21, 25, 32, 38, 43, 48, 49, 56, 63, 68, 73, 81, 88, 94, 99, 107, 113, 119, 120, 121, 125, 126, 127, and 128), mostly due to movement artifacts. The remaining 100 electrodes were re-referenced using their average as reference. Finally, recordings were segmented by trials, taking 25s of each trial (25, 30, 60 or 150 complete cycles for respectively 4, 6, 12 and 30 Hz presentation) starting 2.5s after the beginning of the trial, except for the 4 Hz presentation for which the cropped trials started 3s after the beginning of the actual trial.

For infants’ data, we only considered trials for which we were certain that infants paid attention to the sequence of images. For this, we computed the response at the base frequency (4, 6, 12 or 30 Hz). We applied a Fast Fourier Transform (FFT) to each trial data, over the 3 occipital electrodes (Oz, O1, O2), where we expected to find high visual response to the image streams, provided that infants looked at the stream of images. We averaged the spectra of those 3 electrodes, per trial and per participant, and computed the signal-to-noise ratio (SNR) by dividing the amplitude at the base frequency (i.e., 4, 6, 12, 30 Hz) and 2^nd^ harmonic (i.e., 8, 12, 24, 60 Hz), by the local average signal, that is the mean of amplitudes of the 20 (for 4 Hz presentation rate) or 24 (for 6, 12 and 30 Hz presentation rate) surrounding bins (10 or 12 bins on each side of the amplitude of interest, excluding the immediate adjacent bins). If the SNR at the 1^st^ and/or 2^nd^ harmonic of the image presentation frequency was superior to an arbitrary threshold of 1.5, the trial was included in further analyses.

In addition, we excluded trials for which the experimenter noted online that the infant moved too much, did not look at the screen, touched the mother with their head, pulled off, touched or grabbed the net, or cried. Infants who provided only one trial were not included in the final sample. On average, 4-month-old infants tested with 4 Hz streams contributed 8.06±3.64 trials, 4-month-old infants tested with 6 Hz streams contributed 7.44±4.41 trials, 9-month-old infants tested with 6 Hz streams contributed 7.44±3.56 trials, 9-month-old infants tested with 12 Hz streams contributed 6.72±2.89 trials and 9-month-old infants tested with 30 Hz streams contributed 4.44±2.20 trials.

Included trials were averaged in the time domain for each participant, each electrode and each trial type (animate category-change and inanimate category-change). We then applied a FFT to those averages for examination in the EEG frequency-domain at the high frequency resolution of 0.04 Hz (=1/25 seconds). EEG spectra were averaged across trial types (for adult participants), leading to one EEG spectrum per participant, at each electrode. The baseline-corrected amplitudes were computed for each participant and electrode by subtracting from the amplitude of interest the local baseline, that is the mean of amplitudes of the surrounding bins (10 bins on each side for the 4 Hz presentation; 12 bins on each side for other presentations), excluding the immediate adjacent bins and maximal and minimal bins (*70, 71*). That corresponds to a frequency range of respectively ± 0.44 and ± 0.52 Hz. The number of bins was reduced for the 4 Hz presentation to avoid overlap of bins in computing the baseline-corrected amplitude of 2 different harmonics of the category-change frequency (e.g., 1 and 2 Hz). The baseline-corrected amplitudes were also computed for all bins surrounding each frequency of interest, and used to visualize the data (Figure 3).

### Analysis

#### Group analyses

For each group, we selected the baseline-corrected amplitudes at all harmonics under 50 Hz, excluding harmonics that are common to the visual presentation rate (e.g., 12, 24, 36, 48 Hz for the 12 Hz streams) and summed them (*72*) for each participant, at each electrode. We defined as a priori region of interest the 27 electrodes covering the posterior area of the scalp, where a (visual) response should be recorded (*43*) (electrodes 58, 59, 60, 61, 62, 64, 65, 66, 67, 69, 70, 71, 72, 74, 75, 76, 77, 78, 82, 83, 84, 85, 89, 90, 91, 95, 96). Those electrodes include the Oz, O1, O2, Pz, P7 and P8 electrodes (respectively 75, 70, 83, 62, 58 and 96 in EGI nomenclature) of the international 10-10 system. For each participant, baseline-corrected amplitudes were summed over all harmonics of F_C_ under 50 Hz, excluding harmonics that are common to the visual presentation rate (e.g., 12, 24, 36, 48 Hz for the 12 Hz streams). This sum was averaged across the 27 posterior electrodes. We then performed a one-sample *t-*test one-sided to compare the group response to the noise-level (zero).

Finally, to localize the response more precisely, we performed a cluster mass permutation analysis in these posterior electrodes. For each posterior electrode, we performed a one-sample *t*-test to compare the sum of the baseline-corrected amplitude to the noise-level (zero). Neighboring electrodes showing an above threshold effect (*P* < .05) were clustered and tested for significance using nonparametric cluster-mass permutation test (*73*) with 5000 random sign permutations of the original data.

#### Comparisons between groups

The responses measured in the different groups were also compared across Experiments including participant of the same age. We looked for differences between groups computing the sum of baseline-corrected amplitudes over all harmonics under 50 Hz, averaging over the posterior electrodes for each participant, using the same parameters used in the group analysis. Expecting a larger response for slower presentation rates, we performed a one-sided two-sample *t*-test to compared the two 4-month-old infant groups (Experiments 1-2), and one-way ANOVAs to compare the three 9-month-old infant groups (Experiments 3-5) and the three adult groups (Experiment 6) respectively.

### Decoding

#### Preprocessing

All preprocessing was performed using EEGLAB toolbox (*69*) and MATLAB R2023b (MathWorks). Raw EEG data for each participant in each group were first filtered using Hamming windowed sinc FIR filter at 0.1 Hz (high-pass) and at 100 Hz (low-pass). All electrodes were then re-referenced using the average of all electrodes as a reference. Data were then down sample to 250 Hz. Recordings were segmented by images, the continuous data being segmented in epochs between −100 ms and 1000 ms with respect to the onset of the image, and baseline-corrected by subtracting the mean of the 100 ms interval before the onset of the stimuli from the entire epoch. Only epochs corresponding to images of the standard category were extracted and used in the decoding. Epochs corresponding to the category-change images were not considered.

#### Decoding

We perform a multivariate pattern analysis using the linear discriminant analysis of CoSMoMVPA (*74*), conducting all analyses on a participant-specific basis, for each Experiment. More specifically, we conducted a time-resolved decoding analysis, examining the EEG data from −100 ms to 1000 ms with respect to the image onset, every 4 ms. For each group (4 Hz, 6 Hz, 12 Hz and 30 Hz), each participant, at each time point, we extracted the EEG signal at each electrode for each image, and performed a decoding for each combination possible (320^*^319/2 = 51 040 total), separately per domain (animate and inanimate), as we were no longer interested in the animate-inanimate categorization but by finer-grained representation of the images.

As the number of repetitions per objects may vary (2.83±0.63 repetition on average for 4-month-old infants in Experiment 1 across the two domains, 3.88±0.58 repetition on average for 4-month-old infants in Experiment 2, 3.78±0.55 repetition on average for 9-month-old infants in Experiment 3, 6.51±0.54 repetition on average for 9-month-old infants in Experiment 4, 10.70±0.86 repetition on average for 9-month-old infants in Experiment 5, 7.74±0.62 repetition on average for adults in Experiment 6 at 6 Hz), we first randomly selected a subset of epoch for the image that has the most repetitions to match the number of repetition of the image that has the least, providing at least 4 epochs per images to train and test the model. For infant groups, some infants may not have seen all the 320 images: on average, 4-month-olds in Experiment 1 saw 285.38±35.11 (minimum 209, maximum 319 images), in Experiment 2 4-month-olds saw 303.16±24.33 (minimum 237, maximum 320 images), in Experiment 3 9-month-olds saw 308.75±13.48 (minimum 275, maximum 320 images), in Experiment 4 9-month-olds saw 319.84±0.72 (minimum 316, maximum 320 images) and in Experiment 5, 9-month-olds saw 319.97±0.18 (minimum 319, maximum 320 images). In addition, they do not necessarily have at least 4 epochs per image. Thus, decoding accuracies for those pairs of images that were either not seen or not seen enough were not computed.

With the acceleration of the speed of presentation comes many more presentations of the images (thus many more epochs per images). In order to maintain an equivalent training regime of the model at different frequencies, we randomly selected, for each adult group tested at 12 or 30 Hz in Experiment 6, as many repetitions of the images as those used for the group of 6 Hz. We did not do that for infant groups, as we wanted to maximize the possibility of finding significant decoding between images, unsuccessfully.

We employed a leave-one-out cross-validation approach, dividing the selected epochs into training set and testing set to perform a pairwise decoding between all possible combinations of images. We repeated this operation as many times as needed for each epoch to be in the testing set once. We further repeated this operation 5 times, to have at least 5 random selections of epochs of the most seen image, and averaged the results across those repetitions, to generalize the result at the image-level and not the specific random selection. We then arranged the decoding accuracies for each pair of images into a 320 by 320 decoding accuracy matrix (EEG-RDM) per domain, with rows and columns corresponding to the decoded images. This matrix is symmetric across the diagonal, with the diagonal being undefined.

We performed a cluster-mass permutation analysis on the average of all decoding pairs for each group, averaging the accuracy across domains for adult participants in Experiment 6 as they all saw images of both domains, and considering the 32 infant participants all together in Experiments 1-5 as each infant only saw images of one domain. Every 4 ms, at each time-point, we performed a one-sample one-tailed *t*-test to compare the decoding accuracy to chance-level (0.5). Neighboring time points showing an above threshold effect (*P* < .05) were clustered and tested for significance using nonparametric cluster-mass permutation test (*73*) with 5000 random permutations of signs of the original data. This analysis revealed above chance performance only in Experiment 6, for adult participants. Further analyses were thus only performed for Experiment 6.

We compared groups tested with different speeds of presentation, performing another cluster-mass permutation analysis using a one-way ANOVA at each time-points. Following the cluster-mass analysis comparing all groups together, we performed two-sample *t*-tests to compare each group to the other, in the previously defined time-windows, averaging the decoding accuracy in the time-windows for each group.

### Representational similarity analysis (RSA)

We compared the representation of images, as yielded by the decoding analysis and summarized in the EEG-RDM, to the representation of those same images in Deep Convolutional Neural Networks (CNN). We first considered AlexNet, a CNN that provides a reliable model of the primate visual system, the first layer of the model representing low-level features and later layers higher-level features (*75*).

AlexNet was trained on approximately 1.2 million images from the ImageNet database (*52*), encompassing 1,000 classes including animals (40%) and objects (60%). We used a pretrained version of this model available in MATLAB (MatConvNet), with standardized preprocessing steps such as mean subtraction of the training images and scaling of all stimuli to 224×224 pixels.

We extracted features from convolutional layers (‘conv1’ through ‘conv5’) and fully connected layers (‘fc6’, ‘fc7’, and ‘fc8’). For each layer, for each domain, we computed a dissimilarity matrix (CNN-RDM; *76*) representing the pairwise dissimilarities between the features extracted from each image, computed as 1-*rho*, where *rho* is the *Pearson* coefficient of the correlation between vectors of features extracted for the two images compared.

Performing a Representational Similarity Analysis (RSA; *76*), we computed the *Spearman* correlation between the CNN-RDM and the EEG-RDM at each time point. We first transformed the upper triangle of each matrix, for both domains, into a vector, and concatenated together the two vectors across domains, resulting in a vector of 102080 combinations (51040 per domains) for EEG and for CNN, in order to perform the correlation considering both domains at the same time. We then used sign permutation tests and cluster-mass analyses to determine significant windows showing correlation between the EEG-RDMs and the CNN-RDMs for each group and each layer of the CNN.

We compared groups at each layer by performing a cluster-mass permutation analysis with a one-way ANOVA at each time point. Following the cluster-mass analysis, we performed two-sample *t*-tests to compare each group to the other, in the cluster of time points revealed by the ANOVA, averaging the coefficient of correlation in this cluster for each group.

## Author contributions

Conceptualization: C.S., L.P., J-R.H.; Methodology: C.S., L.P., J-R.H.; Software: C.S., M.A.; Investigation: C.S., E.S., S.N.; Formal analysis: C.S., M.A., L.P., J-R.H.; Funding acquisition: C.S., D.K., L.P., J-R.H.; Supervision: L.P., J-R.H.; Writing – original draft: C.S., J-R.H.; Writing – review & editing: C.S., E.S., S.N., Z.B., S.D., D.K., L.P., J-R.H.

## Data, code, and materials availability

Data, code, materials, and a reproducible manuscript are available at https://osf.io/jsvh4/.

## Supplementary material

- Supplementary results
- Experiment S1
- Video S1: Movie of a trial 4Hz
- Video S2: Movie of a trial 4Hz, phase-scrambled images
- Video S3: Movie of a trial 6 Hz
- Video S4: Movie of a trial 12 Hz
- Video S5: Movie of a trial 30 Hz
- Video S6: dog original
- Video S7: dog - acceleration x3
- Video S8: dog - acceleration x7

## Footnotes

1 Note that a similar analysis of infant data did not yield any significant decoding, probably due to insufficient amount of data (*Supplementary material*)

