## Supplementary Materials for "Acceleration of vision changes object representations from infancy to adulthood"

#### *Supplementary Material*

Céline Spriet[1,2], Emilie Serraille[1,2], Samar Naderi[1,2], Marwane Agrebi[3], Zied Bouyahia[2,3], Stéphane Derrode[2,3], Daniel Kaiser[4,5,6,7], Liuba Papeo[1,2], Jean-Rémy Hochmann[1,2]*

[1]CNRS UMR5229 – Institut des Sciences Cognitives Marc Jeannerod, 67 Bd Pinel 69675 Bron, France

[2]Université Claude Bernard Lyon 1, France

[3]Ecole Centrale de Lyon, LIRIS, CNRS UMR5205, France

[4] Neural Computation Group, Department of Mathematics and Computer Science, Physics, Geography, Justus Liebig University Giessen, Germany

[5] Center for Mind, Brain and Behavior (CMBB), Universities of Giessen, Marburg, and Darmstadt, Germany

[6] Center for Applied Computer Science and Data Science (ZAD), Justus Liebig University Giessen, Germany

[7] Cluster of Excellence "The Adaptive Mind", Universities of Giessen, Marburg, and Darmstadt, Germany

**Supplementary material**

- Supplementary results
- Experiment S1
- Video S1: Movie of a trial 4Hz
- Video S2: Movie of a trial 4Hz, phase-scrambled images
- Video S3: Movie of a trial 6 Hz
- Video S4: Movie of a trial 12 Hz
- Video S5: Movie of a trial 30 Hz
- Video S6: dog original
- Video S7: dog - acceleration x3
- Video S8: dog - acceleration x7

#### Supplementary results

##### Representational similarity analysis (RSA)

We compared the representation of images, as yielded by the decoding analysis and summarized in the EEG-RDM, to the representation of those same images in Deep Convolutional Neural Networks (CNN). In addition to AlexNet, presented in the main text, we carried this analysis with 4 other CNNs: VGG19, cornet-RT, cornet-S and cornet-Z. Results converge with those obtained for AlexNet and are plotted on Figures S1 and S2.

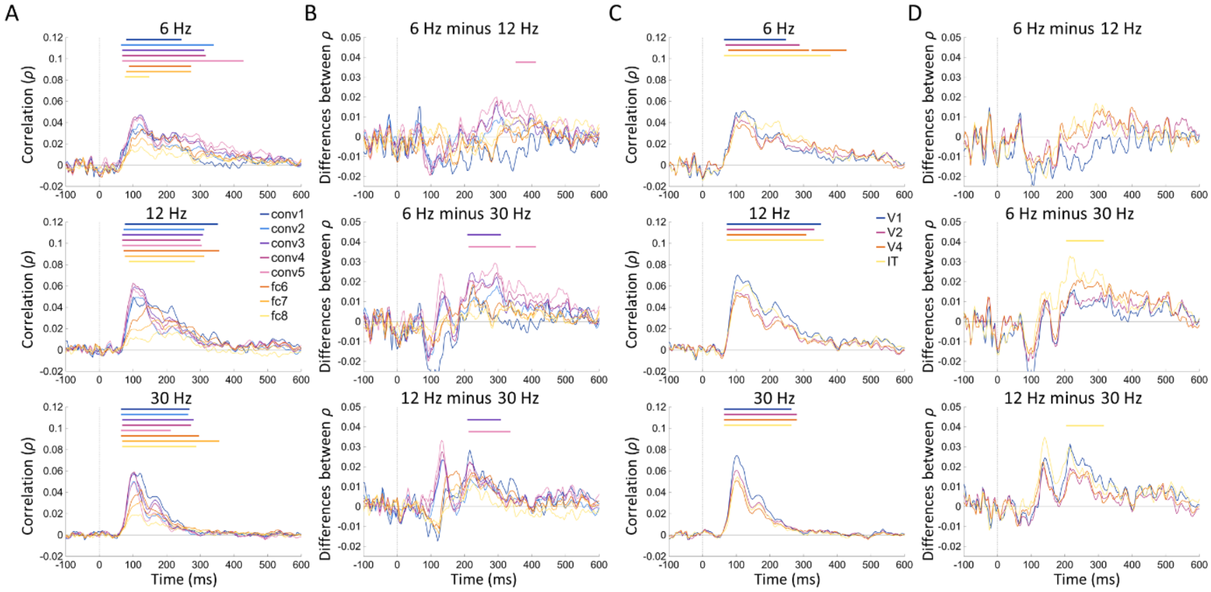


**Figure S1 – RSA with VGG-19 & cornet-RT.** (A) Results of the RSA between the EEG decoding and 8 layers of VGG-19 at each time point of the EEG signal. Significant correlations are denoted by colored bars for each layer. (B) Differences between the correlation of each group and layer of VGG-19. Significant differences between groups are denoted by colored bar for each layer. (C) Results of the RSA between the EEG decoding and the 4 layers of cornet-RT at each time point of the EEG signal. Significant correlations are denoted by colored bars for each layer. (D) Differences between the correlation of each group and layer of cornet-RT. Significant differences between groups are denoted by colored bar for each layer.

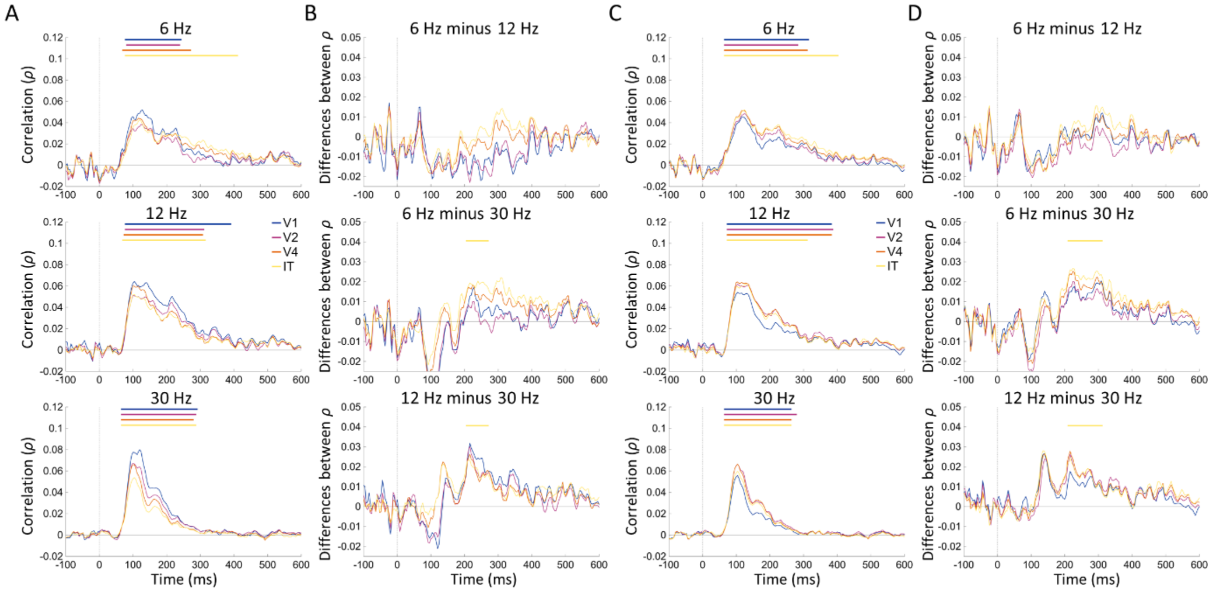


**Figure S2 – RSA with cornet-S & cornet-Z.** (A) Results of the RSA between the EEG decoding and the 4 layers of cornet-S at each time point of the EEG signal. Significant correlations are denoted by colored bars for each layer. (B) Differences between the correlation of each group and layer of cornet-S. Significant differences between groups are denoted by colored bar for each layer. (C) Results of the RSA between the EEG decoding and the 4 layers of cornet-Z at each time point of the EEG signal. Significant correlations are denoted by colored bars for each layer. (D) Differences between the correlation of each group and layer of cornet-Z. Significant differences between groups are denoted by colored bar for each layer.

**Analyses of the Base-frequency response**

In Experiments 1-5, trials were selected for showing a response at the base frequency, so that groups are guaranteed to show a response that differs from chance at the base frequency. Here, we tested whether the value of the base frequency impacted that response across experiments. Results are plotted in Figure S3.

*Experiments 1-2* - The base-frequency response was significantly higher in the group tested with 4 Hz-streams than in the group tested with 6-Hz streams, over posterior electrodes ($M_{Difference}\pm sd$ = 0.217±0.313; 95% CI = 0.086 – Inf; $t(62) = 2.775$; $P = .004$; $d = 0.694$).

*Experiments 3-5* – A one-way ANOVA revealed a significant effect of the value of the base frequency (6, 12, 30-Hz), showing that the base-frequency response over posterior electrodes decreased as the stimulus presentation rate increased ($F(2,95) = 60.285$; $P < .0001$; $\eta^2 = 0.565$). The response was higher for 6-Hz streams *vs.* 12-Hz streams ($M_{Difference}\pm sd$ = 0.415±0.315; 95% CI = 0.283 – Inf; $t(62) = 5.260$; $P < .0001$; $d = 1.315$), for 6-Hz streams *vs.* 30-Hz streams ($M_{Difference}\pm sd$ = 0.704±0.298; 95% CI = 0.580 – Inf; $t(62) = 9.448$; $P < .0001$; $d = 2.362$) and for 12-Hz streams *vs.* 30-Hz streams ($M_{Difference}\pm sd$ = 0.290±0.106; 95% CI = 0.246 – Inf; $t(62) = 10.927$; $P < .0001$; $d = 2.732$).

*Experiment 6* - In adults, the base-frequency response over posterior electrodes also varied with the value of the base frequency (one-way ANOVA: $F(2,35) = 47.002$; $P < .0001$; $\eta^2 = 0.740$). In effect, the base-frequency response was higher for 6-Hz as compare to 12-Hz streams ($M_{Difference}\pm sd$ = 0.853±0.345; 95% CI = 0.611 – Inf; $t(22) = 6.057$; $P < .0001$; $d = 2.473$), and 30-Hz streams ($M_{Difference}\pm sd$ = 1.054±0.336; 95% CI = 0.819 – Inf; $t(22) = 7.689$; $P < .0001$; $d = 3.139$) and higher for 12-Hz streams as compare to 30-Hz streams ($M_{Difference}\pm sd$ = 0.201±0.090; 95% CI = 0.138 – Inf; $t(22) = 5.462$; $P < .0001$; $d = 2.230$).

The base-frequency response was significantly higher than noise-level over posterior electrodes in all adult groups (6 Hz group: $M_{Amplitude}\pm sd$ = 1.148±0.473; 95% CI = 0.903 – Inf; $t(11) = 8.412$; $P < .0001$; $d = 2.428$; 12 Hz group: $M_{Amplitude}\pm sd$ = 0.295±0.120; 95% CI = 0.233 – Inf; $t(11) = 8.541$; $P < .0001$; $d = 2.466$; 30 Hz group: $M_{Amplitude}\pm sd$ = 0.094±0.044; 95% CI = 0.071 – Inf; $t(11) = 7.390$; $P < .0001$; $d = 2.133$).

At all ages, both the category-change response and the base-frequency response decreased when increasing the stimulus presentation rate. We further asked whether the effect observed for the category-change and the effect observed for the base-frequency response followed the same pattern. To this end, we performed a mixed-ANOVA over posterior electrodes, with one between-subject factor Presentation Rate and one within-subject factor Response of Interest (category-change response, base-frequency response). A significant interaction of those factors would show that Presentation Rate affects differently the category-change response and the base-frequency response.

In Experiments 1-2 with 4-month-olds, we found a significant effect of Presentation Rate ($F(1,62) = 9.977$, $P = .003$; $\eta^2 = 0.046$) and a significant effect of Response of Interest ($F(1,62) = 109.228$, $P < .0001$; $\eta^2 = 0.428$), but the interaction between the two factors was not significant ($F(1,62) = 0.123$, $P = .727$; $\eta^2 = 0.0005$).

In Experiments 3-5, with 9-month-olds, we found a significant effect of Presentation Rate ($F(2,93) = 40.839$, $P < .0001$; $\eta^2 = 0.282$), a significant effect of Response of Interest ($F(1,93) = 22.857$, $P < .0001$; $\eta^2 = 0.068$) and a significant interaction between the two factors ($F(2,93) = 8.849$, $P = .0003$; $\eta^2 = 0.053$).

In Experiment 6, with adults, we found a significant effect of Presentation Rate ($F(2,33) = 38.411$, $P < .0001$; $\eta^2 = 0.451$), a significant effect of Response of Interest ($F(1,33) = 22.962$, $P < .0001$; $\eta^2 = 0.067$) and a significant interaction between the two factors ($F(2,33) = 32.991$, $P < .0001$; $\eta^2 = 0.193$).

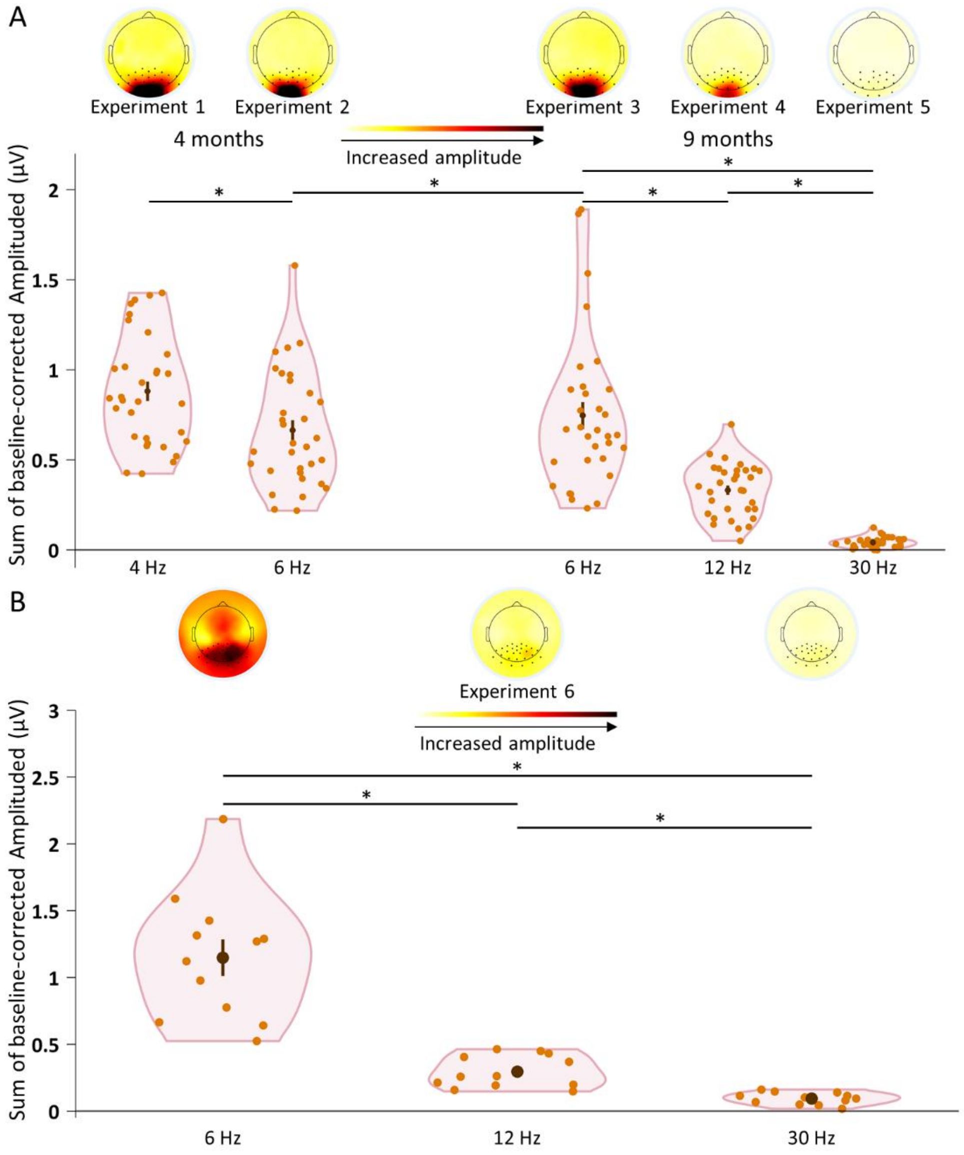


**Figure S3 – Base-frequency response for stimulus streams of different presentation rates and at different ages.** Results of Experiments 1-5 (A) and Experiment 6 (B). Sum of baseline-corrected amplitudes for harmonics under 50 Hz, averaged over posterior electrodes and all participants of each group, showing a difference between groups for each age in addition of a significant response for each group (localized with dark dots in the topoplots). Baseline-corrected amplitude responses were computed by subtracting the average amplitude of surrounding frequencies from the amplitude of the base-frequency response. Orange dots represent the response of each participant; brown dot represent the mean response of the group; lines represent the standard error of the mean.

#### Decoding results

After computing the decoding accuracy, we assessed statistical significance for each group in all Experiments using cluster-mass permutation test at each time point, performing a one-sided *t*-test versus 0 after subtracting chance-level (0.5) from the accuracy. No significant decoding was found for 4-month-olds in Experiments 1-2 nor 9-month-olds in Experiments 3-5. Significant decoding between objects were found for each of the adult groups in Experiment 6, starting earlier for slowest presentation rate, and ending earlier for fastest presentation rate (6 Hz, time-window = [56 588], $M_{Accuracy\%} \pm sd$ = 51.704±0.643, min-max = 50.221-53.614, $P < .001$; 12 Hz, time-window = [68 396], $M_{Accuracy\%} \pm sd$ = 51.224±0.405, min-max = 50.140-52.919, $P < .001$; 30 Hz, time-window = [60 328], $M_{Accuracy\%} \pm sd$ = 50.638±0.256, min-max = 50.052-51.910, $P < .001$).

### Experiment S1

Experiment S1 aimed at showing that the effects observed in Experiments 1-5 are not due to low-level properties of the stimuli. One group of 4-month-old infants was tested with the same procedure as Experiment 1 (base frequency 4 Hz; category-change frequency 1 Hz), but using phase-scrambled images. Phase-scrambled images match the original images for low-level features such as surface area, spatial frequency distribution, and luminance, color and contrast variations, but crucially, they are unrecognizable (video S2).

#### Material and Methods

*Participants:*

Experiment S1 involved 32 4-month-old infants tested with 4 Hz streams (10 males; mean age 137.7±8.3 days). Five additional infants were tested in that Experiment but excluded from the analysis due to an insufficient number of trials included (see preprocessing), or crying and moving too much during the experiment.

*Stimuli:*

Stimuli for Experiment S1 were the phase-scramble version of the stimuli used in Experiment 1. Phase-scramble stimuli were created using the *imscramble* function (http://martin-hebart.de/webpages/code/stimuli.html) in Matlab, randomizing the spatial spectrum of the stimuli. This manipulation preserved the number of pixels, the color, the contrast, the luminance and the power-spectral distribution of the stimuli intact, but render the object identity unrecognizable by impacting higher-level visual features, especially the shape (video S2).

*Experimental procedure:*

Same as Experiment 1 except for the stimuli.

***FT-EEG***

*Preprocessing*

On average, 4-month-old infants tested with 4 Hz streams in Experiment 1 contributed 8.06±3.64 trials, 4-month-old infants tested with 4 Hz streams in Experiment S1 contributed 7.94±3.30 trials, 4-month-old infants tested with 6 Hz streams contributed 7.44±4.41 trials, 9-month-old infants tested with 6 Hz streams contributed 7.44±3.56 trials, 9-month-old infants tested with 12 Hz streams contributed 6.72±2.89 trials and 9-month-old infants tested with 30 Hz streams contributed 4.44±2.20 trials.

#### Results

*Base-frequency response*

As in Experiments 1, trials in Experiment S1 were selected for showing a response to the base-frequency (SNR at 1 or 2 Hz > 1.5). That response was not different in Experiment 1 with intact images and Experiment S1 with phase-scrambled images ($M_{Difference} \pm sd$ = -0.229±0.423; 95% CI = -0.405 – Inf; $t(62)$ = -2.166; $P$ = .983; $d$ = 0.541).

*Category-change response*

A one-sample one-sided $t$-test over posterior electrodes revealed no category-change response ($M_{Amplitude} \pm sd$ = 0.075±0.435; 95% CI = -0.056 – Inf; $t(31)$ = 0.969; $P$ = .170; $d$ = 0.171 and no cluster of electrodes found).

The category-change response over posterior electrodes was significantly higher in Experiment 1 with the original images, than in Experiment S1 with phase-scrambled images (two-sample $t$-test; $M_{Difference} \pm sd$ = 0.183±0.397; 95% CI = 0.017 – Inf; $t(62)$ = 1.841; $P$ = .035; $d$ = 0.460; Figure S4).

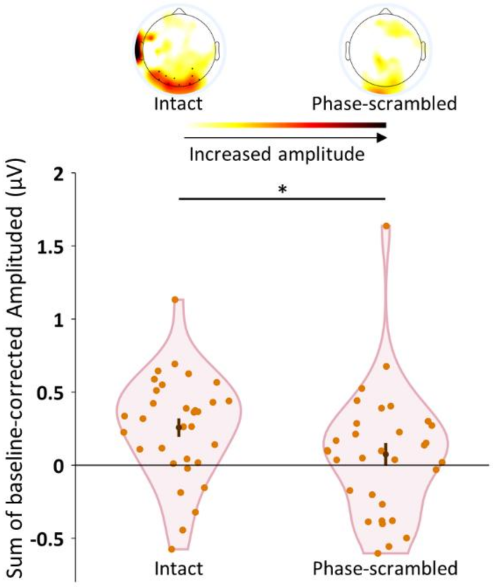


**Figure S4 – Category-change frequency response for 4-Hz streams of intact (Experiment 1) and phase-scrambled images (Experiment S1) at 4 months.** Sum of baseline-corrected amplitudes for harmonics under 50 Hz, averaged over posterior electrodes and all participants of each group, showing a difference between groups. Baseline-corrected amplitude responses were computed by subtracting the average amplitude of surrounding frequencies from the amplitude of the category-change response. Dark dots in the topoplot represent the electrodes showing a significant response at the group-level. Orange dots represent the response of each participant; brown dot represent the mean response of the group; lines represent the standard error from the mean.

**Table S1. Statistical details for the group differences between the average decoding accuracy**

| Difference | mean accuracy (*SD*) | CI (min – max) | *t*(df) | *P* | Cohen's *d* |
|---|---|---|---|---|---|
| 6 Hz - 12 Hz | .010 (*.005*) | .006 – .014 | 4.712 (22) | < .001 | 1.904 |
| 6 Hz - 30 Hz | .015 (*.005*) | .011 – .019 | 7.651 (22) | < .0001 | 3.102 |
| 12 Hz - 30 Hz | .005 (*.002*) | .003 – .007 | 5.750 (22) | < .0001 | 2.409 |

*Notes: The column "mean accuracy" reports the average differences between groups; CI, 95% confidence interval.*

**Table S2. Statistical details for the RSA results between the AlexNet model and the EEG decoding RDMs at each layer, across different presentation rate**

| Speed | N° layer | Time-window (in ms) | mean ρ (*SD*) | *P* |
|---|---|---|---|---|
| 6 Hz | 1 | [80 248] | .020 (*.017*) | .009 |
| | 2 | [64 316] | .028 (*.020*) | .003 |
| | 3 | [64 316] | .029 (*.019*) | < .001 |
| | 4 | [64 412] | .026 (*.018*) | < .001 |
| | 5 | [68 316] | .029 (*.020*) | .001 |
| | 6 | [88 268] | .032 (*.027*) | .011 |
| | 7 | [88 268] | .027 (*.021*) | .011 |
| | 8 | [80 276] | .017 (*.013*) | .009 |
| | | [288 376] | .012 (*.011*) | .032 |
| | | [720 868] | .011 (*.012*) | .015 |
| | | [880 996] | .012 (*.011*) | .019 |
| 12 Hz | 1 | [76 384] | .020 (*.008*) | < .0001 |
| | 2 | [72 304] | .031 (*.009*) | < .001 |
| | 3 | [68 304] | .031 (*.008*) | < .001 |
| | 4 | [68 308] | .032 (*.008*) | < .001 |
| | 5 | [68 308] | .032 (*.009*) | < .0001 |
| | 6 | [72 384] | .027 (*.010*) | < .001 |
| | 7 | [72 316] | .027 (*.010*) | < .001 |
| | 8 | [80 316] | .017 (*.007*) | < .001 |
| 30 Hz | 1 | [68 280] | .019 (*.007*) | < .001 |
| | 2 | [64 266] | .022 (*.009*) | .001 |
| | 3 | [64 264] | .022 (*.008*) | < .001 |
| | 4 | [64 284] | .022 (*.008*) | .001 |
| | 5 | [60 284] | .023 (*.008*) | .002 |
| | 6 | [64 264] | .021 (*.007*) | .001 |
| | 7 | [64 288] | .018 (*.005*) | < .0001 |
| | 8 | [68 284] | .012 (*.004*) | < .0001 |

*Notes: The column "mean ρ" reports the average ρ in each time window.*

**Table S3. Statistical details for the group comparisons of the correlation at each layer of AlexNet**

| Difference | N° layer | Time-window | mean ρ (*SD*) | CI (min – max) | *t*(df) | *P* | Cohen's *d* |
|---|---|---|---|---|---|---|---|
| 6 - 12 Hz | 3 | [204 312] | .008 (*.016*) | -.006 – .022 | 1.240 (22) | .228 | .506 |
| | 4 | [208 312] | .007 (*.018*) | -.008 – .022 | 1.013 (22) | *n.s.* | .411 |
| | 5 | [208 312] | .006 (*.018*) | -.009 – .021 | .789 (22) | *n.s.* | .320 |
| | **8** | **[744 872]** | **.014 (*.010*)** | **.005 – .022** | **3.317 (22)** | **.003** | **1.350** |
| | | **[880 996]** | **.015 (*.010*)** | **.007 – .023** | **3.810 (22)** | **.001** | **1.552** |
| 6 - 30 Hz | **3** | **[204 312]** | **.022 (*.015*)** | **.009 – .035** | **3.500 (22)** | **.002** | **1.431** |
| | **4** | **[208 312]** | **.021 (*.016*)** | **.008 – .035** | **3.218 (22)** | **.004** | **1.313** |
| | **5** | **[208 312]** | **.021 (*.017*)** | **.007 – .035** | **3.059 (22)** | **.006** | **1.248** |
| | **8** | **[744 872]** | **.011 (*.009*)** | **.004 – .018** | **3.151 (22)** | **.005** | **1.279** |
| | | **[880 996]** | **.012 (*.008*)** | **.005 – .019** | **3.479 (22)** | **.002** | **1.429** |
| 12 - 30 Hz | **3** | **[204 312]** | **.014 (*.008*)** | **.007 – .020** | **4.423 (22)** | **< .001** | **1.813** |
| | **4** | **[208 312]** | **.014 (*.008*)** | **.007 – .021** | **4.323 (22)** | **< .001** | **1.775** |
| | **5** | **[208 312]** | **.015 (*.008*)** | **.008 – .022** | **4.599 (22)** | **< .001** | **1.873** |
| | 8 | **[744 872]** | -.003 (*.006*) | -.007 – .003 | -1.030 (22) | *n.s.* | .424 |
| | | **[880 996]** | -.003 (*.006*) | -.008 – .002 | -1.155 (22) | *n.s.* | .468 |

*Notes: time-windows are in ms; mean ρ are the average difference between groups; CI, 95% confidence interval; highlighted in bold are the significant results, α = .05; n.s. = non-significant results with P > .250.*
